# Transiently depleting RNPS1 leads to perdurable changes in alternative splicing

**DOI:** 10.1101/344507

**Authors:** Jérôme Barbier, Alexandre Cloutier, Johanne Toutant, Mathieu Durand, Elvy Lapointe, Philippe Thibault, Benoit Chabot

**Affiliations:** Department of Microbiology and Infectious Diseases, 1 RNomics Platform, Faculty of Medicine and Health Sciences, Université de Sherbrooke, Sherbrooke, Quebec J1E 4K8, Canada

## Abstract

While robust regulatory mechanisms are expected to control the production of splice variants that confer distinct functions, a low level of stochasticity may be tolerated. To investigate stringency of regulation, we followed changes in the splicing of 192 alternative cassette exons after growth of cancer-derived HCT116 cells and embryonic colonocytes. In both cell lines approximately 15% of alternative splicing events changed by more than 10 percentage points over a 42-day period. We then carried out a cycle of transient depletions targeting RNPS1, a splicing regulator implicated in genomic stability. For alternative splicing units not regulated by RNPS1, the level of splicing changes was similar to the stochastic value obtained after normal growth. However, the frequency of perdurable changes was at least twice that value for splicing events regulated by RNPS1. A swap allele assay performed on four RNPS1-responsive units that underwent splicing changes indicated the presence of mutations mediating this effect. Specifically, a T to C mutation in a RNPS1-responsive exon of *ADARB1* confered exon skipping. Our results suggest that fluctuations in the level of a splicing regulator preferentially impact the integrity of genes encoding transcripts that are regulated by this splicing factor to produce perdurable changes in alternative splicing. We discuss the potential implication of this process in human evolution.

## INTRODUCTION

Alternative splicing of pre-messenger RNAs plays a central role in mammals by allowing protein diversification and complex biological function ^1,2^. In humans, nearly all multi-exon genes are alternatively spliced, with the brain being the most diligent user of this process ^3^^−^^5^. Alternative splicing is often regulated in a tissue-, cell- and stress-specific manner ^1^. Control of alternative splicing based on daily changes in body temperature also occurs ^6^. Splicing control requires the interplay of RNA binding proteins (RBPs) that interact with the pre-mRNA to modulate the recognition of splicing signals, thereby altering steps of spliceosome assembly that define the introns to be removed ^7^. SR and hnRNP proteins are well-known splicing regulators, but dozens of other RBPs have been implicated in splicing regulation ^8^^−^^10^. The rate of transcription elongation and the identity of splicing factors associating with promoters and chromatin components also impact splicing decisions ^10,11^. While robust regulation is expected for splicing events that produce variants of different functions (e.g. secreted or membrane-associated proteins, and pro-death or pro-life apoptotic variants), the production of splice variants displaying minimal functional differences may be more frequently subjected to stochastic variations.

Genomic instability refers to a range of genetic alterations from point mutations to chromosome rearrangements. Genomic instability can be a normal process, for example to diversify antibody production, but it can be deleterious when it promotes errors that overwhelm the DNA repair machinery ^12^. One way to trigger genomic instability is through the depletion of specific RBPs. The current view is that portions of transcribed RNA sequences that are not appropriately bound by RBPs will hybridize back onto the melted DNA immediately upstream of the elongating polymerase. The resulting structure, known as a R-loop, may slow down transcription, promote mutations and hyper-recombination ^13^^−^^15^. mRNA processing factors play a widespread role in preventing R-loop-mediated DNA damage in mammalian cells ^16^. Specifically, splicing factors such as SR and SR-like proteins are important to maintain genomic stability ^17^^−^^19^. Inactivation of SRSF1 provokes the accumulation of R-loops, and overexpression of the SR-related protein RNPS1 can rescue defects caused by the loss of SRSF1 ^18^. Based on these observations, we hypothesized that genomic instability triggered by the depletion of a specific RBP could produce mutations with a perdurable effect on splicing. Our study provides a first test of this model; we show that transient drops in RNPS1 can lead to perdurable changes preferentially affecting the splicing of alternative exons that are normally regulated by RNPS1. These results raise the possibility that physiological, environmental or stress-induced changes in the abundance of a specific RBPs may permanently impact the alternative splicing of events that are normally regulated by these RBPs.

## RESULTS

### Changes in alternative splicing during cell growth in culture

To test the hypothesis that the depletion of a RBP can promote permanent change in splicing we focused on the splicing factor RNPS1 because its overexpression rescues the genomic instability induced by depleting SRSF1 ^18^. A doxycycline inducible plasmid that expresses a shRNA targeting RNPS1 was transfected into HCT116 and three individual sublines were isolated. The expression of RNPS1 after shRNA induction (t7si) was measured, as well as 7 days after removing induction (t7si+7) (**Supplementary Figure 1A**). Quantitative RT-PCR indicates that expression of RNPS1 transcripts dropped by more than 50% in all three sublines, and that expression was back to normal levels 7 days after stopping shRNA induction. Although we could not find a reliable source of commercial anti-RNPS1 antibody to confirm these results, this level of depletion of RNPS1 transcripts in the t7si samples was sufficient to change the alternative splicing of two known targets of RNPS1, *Bcl-x* and *SLIT2* ^20^ (**Supplementary Figure 1B**).

Before proceeding with testing our hypothesis, we first wished to ascertain the extent by which alternative splicing profiles varied in HCT116 cells during normal growth in culture. For this we used endpoint RT-PCR to follow 192 alternative splicing events (ASEs) representing cassette exons. We compared profiles between t0 and t42 in culture for the three sublines in the absence of doxycycline in the media. A change was attributed when the difference in Percent Splicing Index (ΔPSI = PSI_t42_ – PSI_t0_) was larger than 10 percentage points (Z-score of 1.23 calculated for biological triplicates on 2137 ASEs with a median variation of 1.83 percentage points). The average frequency of change was 12.8% ± 8.7%, with 61 of the 192 ASEs producing a change in at least one subline (**Supplementary Figure 2**). No ASE changed in all three sublines; 13 ASEs changed in two sublines, with a different polarity seen in three cases. For PSI changes larger than 15 percentage points, the average frequency of change dropped to 5.0% ± 3.9%, with 25 of the 192 ASEs changing in at least one subline, and 4 ASEs changing in 2 sublines (**Supplementary Table IA**). We also recorded the appearance and disappearance of unassigned amplicons relative to t0, with the caveat that the identity of these products was not validated by sequencing. When all types of changes were considered (i.e. ΔPSI larger than 10 percentage points on annotated events, and appearance/disappearance of unassigned amplicons), 28.5% ± 8.0% of the ASEs displayed a change at t42, with 102 of the 192 ASEs experiencing a change when all three sublines are considered (**Supplementary Table IB**).

The average rate of change at shorter time points (7 and 14 days) was equivalent. Using a subset of 96 ASEs, the average frequency of ΔPSI larger than 10 percentage points was 18.4% ± 6.3%, 17.0% ± 9.5% and 10.1% ± 5.7% for t7, t14 and t42, respectively (**Supplementary Table IIA)**. Depending on the subline, between 45% and 100% of the changes detected at a given time point disappeared or changed polarity at a following time point. When all types of changes were considered, the average rate of changes was 30.9% ± 9.1%, 24.6% ± 11.4% and 21.5% ± 4.3% for t7, t14 and t42, respectively (**Supplementary Table IIB**). These results suggest that a mixture of events may be occurring to drive these changes, such as stochastic and stable variations in the expression of splicing regulators, or mutations at splicing signals or regulatory elements.

HCT116 cells have a defective repair machinery ^21^ that may increase genomic instability, which in turn may contribute to changing splicing profiles. To address this question, we tested a normal human embryonic epithelial intestinal cell line (HIEC) ^22^. Three HIEC sublines were grown for 14 days. Based on cell doubling times, HIEC and HCT116 cells underwent a similar number of cell divisions during that period (12.5 and 12.0, respectively). The average frequency of splicing change (annotated and unassigned amplicons) for the 96 ASEs tested was 25.3% ± 2.3% for HIEC (**Supplementary Table III**). A similar frequency of splicing change between HCT116 cancer sublines and normal colonocytes suggests that cancer-associated genomic instability is unlikely to be the main cause of splicing change in this assay.

### Perdurable changes in splicing caused by the transient depletion of RNPS1

Having established the background rate of changes in alternative splicing, we then tested the hypothesis that the transient depletion of a specific splicing regulator could affect splicing profiles in a perdurable way. To identify ASEs that are normally regulated by RNPS1, we induced shRNPS1 expression in our three HCT116 sublines (doxycycline treatment for 7 days) and tested the alternative splicing of 192 ASEs. A reactive unit was defined for each subline by a change of at least 10 percentage points. An average of 24.5% ± 5.0% of the ASEs responded to the depletion of RNPS1 (**Supplementary Table IVA**), with 17 ASEs affected in all three sublines (*BTC, PALM, DMBX1, ITGA6, APRT, PP1L2, PPP1CB, SYNE2, NFAT5-2, DUSP6, UEVLD, DGLUCY-1, COL13A1, C1orf43, PITPNC1, SLC35B2* and *ZMYND8-2*), 24 ASEs affected in 2 sublines, and 28, 6 and 8 ASEs uniquely affected in sublines 1, 2 and 3, respectively. Five ASEs reacted with different polarity in different sublines (*DMBX1, PAXBP1, SNRPB2*, *NFAT5-1* and *ENO3*), and hence were not further considered in the study.

RNPS1 is also an auxiliary component of the exon junction complex (EJC) and participates in non-sense mediate RNA decay (NMD). We asked if the profile of our set of RNPS1-responsive ASEs was controlled by NMD. Two sublines were treated with the translation inhibitor emetine, which blocks translation and hence NMD ^23^. The production of amplicons for three ASEs was affected by emetine, but only *NFAT5-1* produced a PSI shift in the same direction as ΔRNPS1, and in only one of the two sublines tested (**Supplementary Table IVB**), indicating that NMD is not appreciably modulating the profile of splice variants in our set of selected events.

We then performed successive rounds of induction of shRNPS1 in the three HCT116 sublines (three 7-day inductions each followed by a 7-day recovery period). Considering ΔPSI larger than 10 percentage points, an average of 37.6% ± 2.1% of the RNPS1 responsive (RNPS1^R^) ASEs had shifted in the three sublines, while 14.7% ± 7.1% of the RNPS1 nonresponsive (RNPS1^NR^) ASEs had changed (**Figure 1; Supplementary Table IVC and IVD; Supplementary Figure 3**). The difference in the frequency of change between RNPS1^R^ and RNPS1^NR^ ASEs was statistically significant (*P* value = 0.01 using Student t-test). Many of the shifts seen in one or in two sublines also occurred but to a lesser extent in the remaining sublines (i.e. with shifts of 5 to 9 percentage points in *AGAP1, MEIS2, ADARB1-2, HNRNPH3, CPPED1, KTN1-2, ARFIP-1, BAZ1A, ZMIZ2, MBD1, MCL1, HNRNPC, TRAPPC6B*). If we consider that a RNPS1^R^ ASE in one cell subline should be viewed as a RNPS1^R^ ASE in all sublines, 22 changes can be added to the RNPS1^R^ group (black boxes in **Figure 1**) leading to a total of 74 perdurable shifts in 78 RNPS1^R^ ASEs in the three sublines (31.6% of all 234 possible changes). In contrast, 12.5% of the consistent RNPS1^NR^ ASEs had splicing changes (41 out of 327 possible changes) (*P* value for the difference in shifts between RNPS1^R^ and RNPS1^NR^ ASEs = 0.02). If we remove from this compilation the 45 ASEs that had a greater propensity to shift upon growth (i.e. ASEs that shifted in at least two sublines or two time points based on Supplementary Figure 2 and Supplementary Table IA), the difference in perdurable changes between RNPS1^R^ (46 changes out of 126 = 36.5%) and RNPS1^NR^ (32 changes out of 300 = 10.7%) ASEs remains significant (*P* value = 0.04).

**Figure 1.**
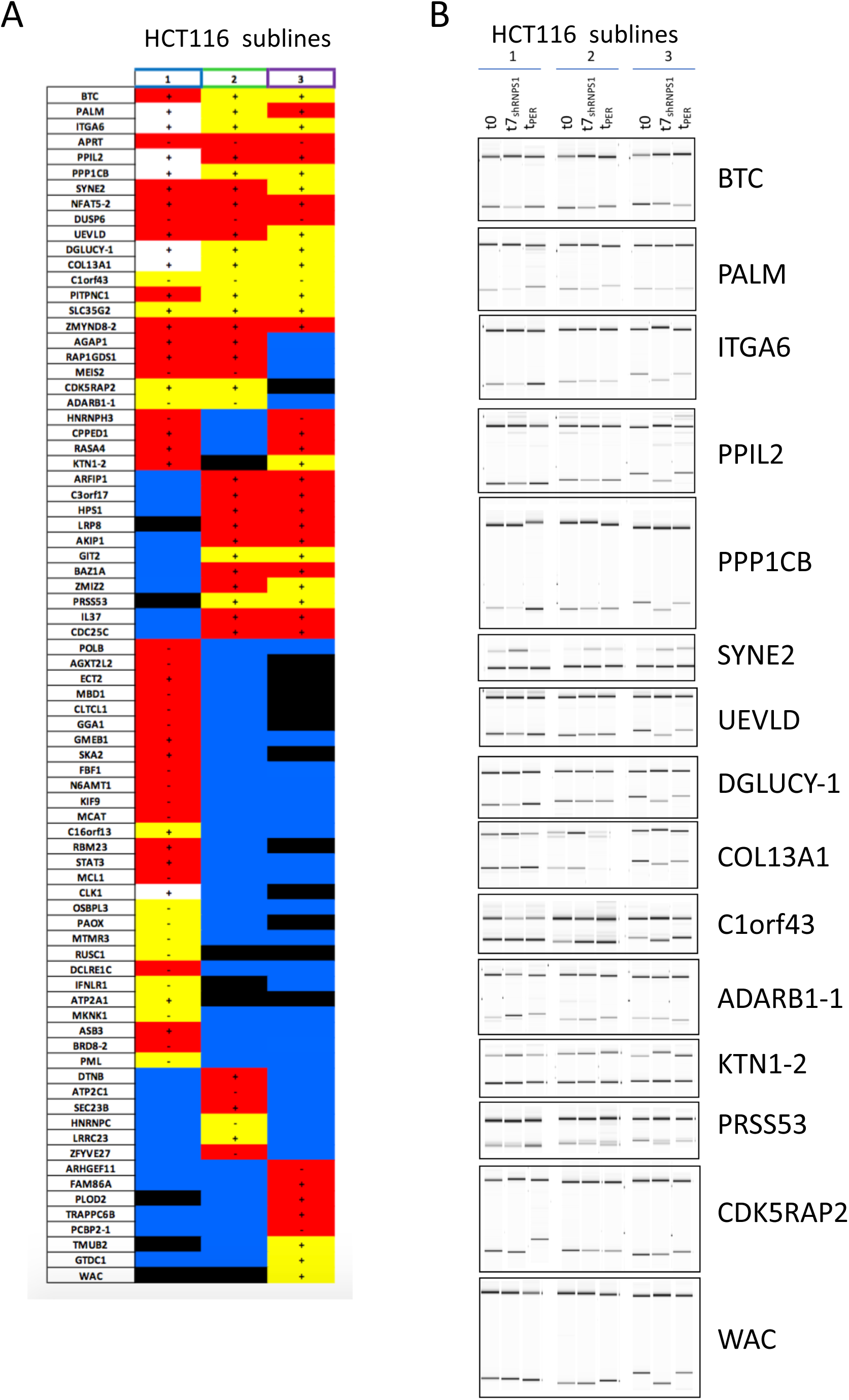
**A.** Perdurable changes in splicing after 3 rounds of transient RNPS1 depletion. Our 78 ΔRNPS1^R^ units are listed with the original impact of depleting RNPS1 indicated by a + (more exon inclusion) or a – (more exon skipping). Yellow boxes indicate a perdurable change occurring in the same polarity as the direct effect of depleting RNPS1. White boxes indicate a change in polarity relative to the direction of the ΔRNPS1-mediated change. Red boxes indicate RNPS1 responsive units that did not yield a perdurable change. Black boxes indicate a perdurable change not matched by a direct ΔRNPS1-mediated change in that subline. Blue color indicates no change when RNPS1 was depleted. **B**. Gel-like representation of electropherograms showing the splicing profiles of 15 ASEs in three HCT116 sublines at t0, in cells induced to express shRNPS1 for 7 days (t7_shRNPS1_), and the perdurable splicing change noted after three rounds of depletion and 7 days of final recovery (t_PER_). Components of sets were often fractionated on different Caliper runs explaining imperfect co-migration of bands. Original electropherograms are provided in Supplementary Figure 3 and at https://rnomics.med.usherbrooke.ca:8080/palace/data/related/2914.

Given that cells had a full week to recover after the last depletion of RNPS1, it is unlikely that shifts were caused by a delayed recovery in the splicing profiles of RNPS1^R^ units. If it had been that case, a greater level of systematic effect would have been expected. Morever, 7 of the 20 perdurable changes in subline 1 occurred in a polarity different from the impact of a direct depletion of RNPS1 (white boxes in **Figure 1**).

### Mutations in ASEs cause perdurable splicing changes

The increased frequency of change in RNPS1-responsive ASEs could be due to *cis*-acting or *trans*-acting mechanisms. A *trans*-acting mechanism would occur if an alteration in the expression or activity of a regulatory RBP (RNPS1 or another splicing regulator). In contrast, a *cis*-acting effect could be due to a mutation in the alternative splicing unit itself that may have been triggerred by the transient depletion of RNPS1. Consistent with the view that the depletion of RNPS1 can trigger R-loop-mediated genomic instability, the RNPS1 depletion stimulated the phosphorylation of H2AX, a known marker of genomic instability (**Supplementary Figure 4**). Co-expression of RNase H1, which reduces the accumulation of R-loops, prevented this stimulation (**Supplementary Figure 4)**.

To distinguish between *trans*- and *cis*-acting mechanisms, we conducted a swap-allele assay. The assay involves amplifying the genomic copy of an ASE in t0 cells and in cells that have experienced the transient RNPS1 depletion protocol (t42_PER_). Units were cloned in minigenes that were then transfected in t0 and t42_PER_ cells (**Figure 2A**). If the change in splicing is due to a mutation, the t42_PER_ unit should be spliced the same way in t0 and t42_PER_ cells. If the effect is *trans*-acting, the t42_PER_ unit should be spliced differently in t0 and t42_PER_ cells, and the t0 unit should be spliced like a t42_PER_ unit in t42_RNPS1_ cells. Units of *ADARB1-1* were amplified, cloned and transfected using the above protocol. The t0 unit of *ADARB1-1* was spliced similarly in t0 and t42_PER_ cells (boxes 1 and 3 in **Figure 2B**, left panel). The splicing profile of the t42_PER_ unit of *ADARB1-1* transfected in t0 and t42_PER_ cells was also consistent with a *cis*-acting mechanism (boxes 2 and 4; **Figure 2B**, left panel). t0 and t42_PER_ units of *PPIL2*, *ABCD4* and *IFNLR1* transfected in t0 cells produced splicing profiles also consistent with the existence of mutations in their t42_PER_ units (boxes 1 and 2 in **Figure 2B**, right panel).

**Figure 2.**
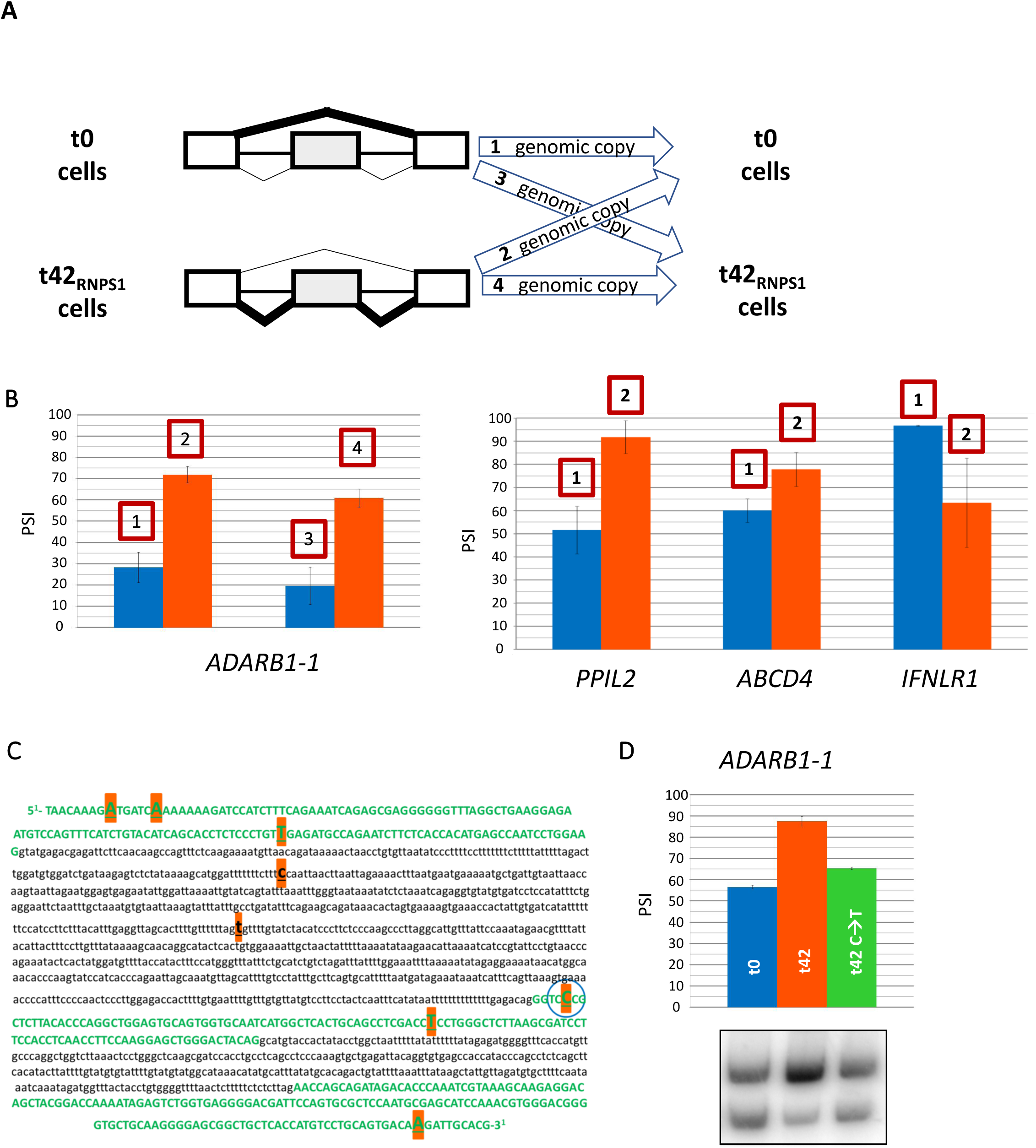
The depletion of RNPS1 produced mutations that affect alternative splicing. **A.** Swap allele assay. Minigenes were constructed by amplifying and cloning genomic DNA from t0 and t42_PER_ cells for units displaying altered splicing profiles. **B.** RT-PCR assays on total RNA from t0 and t42PER cells transfected with plasmid containing the *ADARB1-1* minigenes from t0 and t42_PER_ cells. The boxed numbers refer to the type of allele swap shown in panel A. RT-PCR assays from t0 cells transfected with plasmids containing splicing units was also performed for the three indicated genes. Plasmid-specific primers were used. **C.** Sequence of the alternative unit of *ADARB1-1*. The positions that differ between the t0 and the t42_PER_ units are indicated. Exon sequence is in red capital letters, intron sequence is in black lower case letters. Mutations in t42 relative to the t0 sample are indicated by red boxes. The nucleotide converted into C from the original G in the t0 sample is circled. **D.** RT-PCR analysis from HCT116 cells transfected with plasmids containing the *ADARB1-1* unit from t0 and t42_PER_, as well as the corrected t42_PER_ C➔T version.

The cloned t0 and t42_PER_ genomic units of *ADARB1-1* were sequenced to reveal several differences distributed along the ASE **(Figure 2C**). Noting a T to C conversion in the alternative exon of the t42_PER_ *ADARB1-1* unit near the 3’ splice site, we produced a version corrected for a T at this position. This modification restored the t0 splicing profile (**Figure 2D**), indicating that the T to C mutation in the t42_PER_ *ADARB1-1* unit strongly contributed to the perdurable splicing change.

## DISCUSSION

Although alternative splicing produce proteins with different functions, not all splice variants may be functionally relevant in a specific cellular system. Hence, as long as variants exert no negative impact, stochastic variations in their production may become useful for cell adaptation or as a testing ground for molecular evolution. The number of alternative splicing events that are subjected to random fluctuations is unclear. In the HCT116 colorectal cancer cell line grown for 7, 14 or 42 days, the baseline frequency of variations in the alternative splicing of cassette exons was approximately 10-15% for events displaying variations in PSI greater than 10 percentage points. Although HCT116 cells have a defective DNA repair machinery ^21^, the same rate of splicing changes was observed in the normal fetal colonocyte cell line HIEC. It remains unclear if genomic instability, externally induced or intrinsic variations in the expression of splicing factors might account for these fluctuations.

Recent work have suggested that drops in the levels of specific RBPs can trigger transcription-associated R-loop formation ^15^. This process has been proposed as a potential mechanism underlying the genomic instability caused by the depletion of SRSF1, and can be mitigated by overexpressing RNPS1 ^17,18^. Based on this work, we postulated that the depletion of RNPS1 may lead to R-loops encompassing regulatory elements, and nearby sequences including adjacent splicing signals Mutations in such regulatory-rich regions could alter alternative splicing in a perdurable manner after the transiently depleted regulator is back to normal level. The results of a cyclic and transient depletion of RNPS1 are consistent with this model: compared to RNPS1-nonresponsive units, RNPS1-responsive units were at least twice as likely to display changes in their splicing profile. A swap allele assay conducted on 4 units suggested that these changes were caused by *cis*-acting events. We further identified a point mutation in the evolved *ADARB1* allele that, when converted to its original nucleotide, restored the original splicing profile. Overall, our results suggest that mutations occurring in transiently RNPS1-depleted cells can lead to perdurable splicing changes that will preferentially affect splicing units normally regulated by RNPS1.

Our finding has wide-ranging implications, as it implies that stable changes in alternative splicing can be accelerated by variations in the levels of specific RBPs. These changes may occur in somatic tissues during the lifetime of an organism and may contribute to cell adaptation or disease. This process may also be relevant to our understanding of how alternative splicing may have shaped the evolution of mammals. While the evolution of alternative splicing across different evolutionary lineages remains speculative ^24^, it is indisputable that genomic changes that impact alternative splicing profiles in complex organs such as the human brain need to occur first in germ cells. Interestingly, gene expression in the brain is often reproduced in testis ^25^. Thus, transcription-coupled mutagenesis in testis could potentially lead to alternative splicing changes in the brain, and such changes may have sparked, and may still mold, brain evolution. Notably, Rbm5, Sam68, T-STAR, Celf-1 and Ptbp2 play roles in alternative splicing during mammalian spermatogenesis ^26^^−^^29^. Ptbp2 controls over 25% of alternative splicing events during mouse spermatogenesis with a peak of expression that is associated with splicing repression of transcripts encoding splicing regulators such as SRSF1, SRSF2, RNPS1, THOC1, as well as RNASEH2A ^30^. Thus, variations in the levels of Ptbp2 could in principle impact the expression of several proteins that contribute to genome stability. Another example is the SR protein RSRC1 that affects intellect and behavior through alternative splicing, and it is expressed both in neurons and testis ^25,31^. Interestingly, small changes in body temperature drive rhythmic SR protein phosphorylation to control alternative splicing ^6^. Since testis are subjected to diurnal variation in temperature ^32^, this may alter the activity of splicing regulators with a potential effect on genomic integrity. The high rate of transcription during mammalian germ cell differentiation also creates a high demand for splicing factors ^33^ and may be conducive to R-loop formation ^34^. Small and transient drops in the expression of specific RBPs may therefore offer a potential mechanism that drives the rapid evolution of splicing units. If this process occurs in brain-relevant alternatively spliced regions that are intrinsically unstable (like coding VNTR regions ^35^), the rate of evolution of such units may even be faster. Overall, this process may help explain the rapid evolutionary increase in complexity of the human brain, as well as provide a mechanism that continues to produce new configurations at a rapid pace.

## MATERIALS AND METHODS

### Minigenes

For the shRNPS1/pTER+ plasmid, oligonucleotide pairs were hybridized and inserted in plasmid pTer + cut with BglII and HindIII. Sequences of the oligonucleotides were: shRNPS1Fwd 5’-GATCCCCAAAGGAAGACCAGTAGGAAATTCAAGAGATTTCCTACTGGTCTTC CTTTGTTTTTGGAAAA-3’ shRNPS1Rev 5’-AGCTTTTCCAAAAACAAAGGAAGACCAGTAGGAAATCTCTTGAATTTCCTA CTGGTCTTCCTTTGGG-3’ Plasmid pcDNA4 t0/neo with FLAG-RNAse H1 was made from a RNAse H1 cDNA produced by reverse transcription/amplification from HCT116 total RNA. The full cDNA was first inserted at the HindIII/BamHI sites in p3XFLAG-CMV. The tagged RNAse H1 was then amplified by PCR and inserted into pcDNA4 t0/neo (HindIII/EcoRI) under the control of the inducible promoter. The PCR products used to create the ADARB1 minigene were synthesized from the genomic DNA of t0 and t42 cells. The reaction conditions were as follows: 5 µl of gDNA (2.5 ng/µl), 12.5 µl of GoTaq Mix (Promega, with a high fidelity polymerase), 0.5 µl forward primer (10 µM), 0.5 µl of reverse primer (10 µM) and 6.25 µl of water. Amplification protocol was as follows: 3 minutes at 95°C, followed by 35 cycles at 95°C for 30 seconds, 55°C for 30 seconds and 72°C for 1 minute, ending with an extension/amplification phase of 10 minutes at 72°C. The amplification products were inserted into plasmid pcDNA3.1+. Plasmids and PCR amplification products were digested with restriction endonucleases and then purified on agarose gel and dephosphorylated with 5U Antarctic phosphatase (NEB) in ANT IX buffer (NEB). Ligation was carried out overnight at 40°C with T4 DNA ligase, buffer OPA IX, 1.0 µl of PEG and 12.5 mM of rATP for a final volume of 10 µl. Sequencing of all plasmids was performed prior to transfection in HCT116 cells.

### Cells

HCT116 cells were grown at 37°C in McCoy (Wisent) medium supplemented with 10% FBS. HIEC cells (kindly provided by Jean-François Beaulieu) cells were grown at 37°C in Opti-MEM medium (Wisent) supplemented with 5% FBS and 10 nM GlutaMAX (Gibco).

For transfection, 600 000 HCT116 cells were seeded in 35 mm^2^ wells. Plasmids pTer+ (selection with blasticidin) and pcDNA4 t0/neo (selection with G418) were used to generate HCT116 cell lines expressing the shRNA targeting RNPS1 and FLAG-RNAse H1, respectively Addition of doxycycline (BD BioSciences) lifts repression allowing expression of the shRNA and RNAse H1. The Tet repressor was expressed from pcDNA6/TR (selection with zeomycin). After 24 hours, 2 µg of plasmid DNA and 5 µl of Lipofectamine 2000 (Invitrogen) were incubated for 20 minutes in a final volume of 100 µl of Opti-MEM before being added to the wells containing 1 ml of McCoy’s medium. Six hours later, the media ws replaced with fresh and complete McCoy medium (10% FBS). Selection was applied 24 hours post-transfection. For the shRNPS1 line we used blasticidin (18 µg/ml; Wisent) and zeocin (400 µg/ml; Invitrogen). For the RNAse H1 line (which was created from a subclone of the HCT116 shRNPS1 line), we used G418 (3 mg/ml; Wisent), blasticidin (9 µg/ml) and zeocin (200 µg/ml). Individual colonies of surviving cells were recovered and resuspended to generate subclones. After selection, drug treatment was maintained at 9 µg/ml blasticidin, 200 µg/ml zeocin (and 1.5 mg/ml G418 for the RNAse H1 line).

For wave of depletions with shRNPS1, t0 was defined as the beginning of the shRNA induction with doxycycline. After 7 days of induction, doxycycline was withdrawn for 7 days. This cycle was repeated two more times for a total of 42 days (t42). To verify the impact of NMD inactivation, 3 µM emetine (Sigma-Aldrich) was added to the culture medium. For the calculation of the population doubling, we used the following formula: PD = Log (Nf / No) / Log2 where No is the quantity of cells initially seeded in the well and Nf is the quantity of cells present in the well at the time of measurement and calculated using a hemacymeter.

### RNA extraction and RT-PCR analysis

Cells were washed with PBS and RNA was extracted using TRIzol (Invitrogen). The RNA pellets were washed twice with 900 µl of cold 70% ethanol and resuspended in 20 µl of Nanopure water (Qiagen). The total RNA extracts were subsequently treated with 1U of DNase I (Fermentas) for 30 minutes at 37°C. DNase I was inactivated by adding EDTA and incubating at 65°C for 10 minutes.

Reverse transcription was performed with random hexamers for the endogenous mRNAs of Bcl-x, SLIT2 and primer RT3 (5’-GAAGGCACAGTCGAGGCTG-3’) for mRNAs produced from minigenes. The RT reaction was carried out at 37°C for 1 hour under the following conditions: 2 µl of RNA, 1.42 µl of water, 0.5 µl of Omniscript 10X buffer (Qiagen), 0.5 µl of dNTPs, 5mM), 0.08 µl of RNase Out (Roche), 0.25 µl of primer (10 µM) and 0.25 µl of Omniscript reverse transcriptase (Qiagen) for a total volume of 5 µl. Sequences of primers for amplification: Bcl-x-X3: 5’-ATGGCAGCAGTAAAGCAAGCG-3’; Bcl-x-X2: 5’-TCATTTCCGACTGAAGAGTGA-3’; SLIT2.F36: 5’-GGCAAGTTTCAACCATATGCC-3’; SLIT2.R4: 5’-GGAGCCATAAATGACTGGTGAC-3’; T7: 5’-TAATACGACTCACTATAGGG-3’; RT4: 5’-CTGATCAGCGGGTTTAAACG-3’

For conventional PCR, 1 µl of cDNA was incubated with 2.5 µl of 10× Taq buffer (NEB), 0.3 µl of dNTP (10 mM), 0.5 µl of each primer (50 µM), 2 µl of Taq polymerase (1U) (NEB) in a total volume of 25 µl. For the radioactive PCR, 0.1 µl of α-^32^P dCTP (PerkinElmer Canada Inc.) was added to the PCR mix. A first cycle of 3 minutes at 95°C was followed by 35 cycles of 30 seconds at 94°C, 30 seconds at 55°C and 30 seconds at 72°C. The PCR reaction was terminated with an extension step of 10 minutes at 72°C. The amplification products were separated in a native gel of 4% polyacrylamide and visualized by autoradiography with a STORM PhosphorImager 860 (GE Healthcare). For alternative splicing profile analysis of the 192 ASEs, RT-PCR reactions were carried out by the RNomics platform of the Université de Sherbrooke. The list of primers used is given in Supplementary Table 1. Reverse transcription was performed using 10 units of Transcriptor reverse transcriptase (Roche), 20 units of RNAseOUT (Invitrogen), 3.2 µg of random hexamers, 1 µM of dNTPs mix, 1X of Transcriptor RT reaction buffer, and 0.2 to 2 µg of total RNA. PCR was performed using 0.2 units of Platinium Taq polymerase, 0.6 µM primers, 1.5 mM MgCl_2_, 10 ng of complementary DNA, 1X PCR reaction buffer and 200 µM dNTP mix. PCR reactions were performed in GeneAmp PCR System 9700 thermocyclers (Thermo Scientific-Invitrogen). A first cycle of 15 minutes at 95°C was followed by 35 cycles of 30 seconds at 94°C, 30 seconds at 55°C and 1 minute at 72°C. PCR reactions were terminated with an extension step of 10 minutes at 72°C. The analysis of amplification products was carried out by capillary electrophoresis with the Labchip GX Touch HT (Perkin Elmer). The percent splicing index (PSI) was calculated for each sample, and ΔPSI (PSI_experimental_ - PSI_control_) was used to monitor the change in splicing. All the results of the endpoint RT-PCR assays on electropherograms can be consulted at https://rnomics.med.usherbrooke.ca:8080/palace/data/related/2914.

### Knockdown analysis

The efficiency of depletion was verified by qRT-PCR analysis using the following protocol: 5 µl of 2X FastStart Universal SYBR Green Master mix (Roche Diagnostics), 10 ng (3 µl) of cDNA, and 200 nM (2 µl) of diluted primer pair solutions for a total volume of 10 µl. The amplitude conditions were as follows: 10 minutes at 95°C followed by 50 cycles of 15 seconds at 95°C, 30 seconds at 60°C and 30 seconds at 72°C. The relative expression level was calculated using the qBASE software. The housekeeping gene used for normalization was PSMC4. Sequence of primers were: PSMC4_global_for_1: 5’-GGCATGGACATCCAGAAG-3’; PSMC4_global_rev_1: 5’-CCACGACCCGGATGAAT-3’; RNPS1 for: 5’-AGGCTATGCGTACGTAGAGTTTG-3’; RNPS1 rev: 5’-GATCTCCTGGCCATCAATTT-3’.

Total proteins were extracted from cells and immunoblot detection was performed with the Western Lighting-ECL (PerkinElmer) chemiluminescence products according to the supplier’s instructions. Primary antibodies were used aas follows: anti-FLAG: mouse, monoclonal clone M2 (Sigma-Aldrich), dilution 1: 1000; anti-α-tubulin: rabbit (Abcam), dilution 1: 3500; anti-H2AX (serine 139): mouse, clone JBW301 monoclonal (Millipore), dilution 1: 1000. Secondary antibodies were either anti-rabbit polyclonals (Cell Signalling 7074) or anti-mouse (BioCan 115-035-003).

## ACKNOWLEDGEMENTS

We thank Jean-François Beaulieu for providing HIEC cells. This study was supported by CIHR grant MOP-10507, the Canada Research Chair in Functional Genomics and a NSERC Discovery grant to BC. BC is the Pierre C. Fournier Chair in Functional Genomics. AC was supported by a NSERC scholarship.

## REFERENCES

1. Baralle, F.E. & Giudice, J. Alternative splicing as a regulator of development and tissue identity. Nat Rev Mol Cell Biol 18, 437–451 (2017).

2. Kalsotra, A. & Cooper, T.A. Functional consequences of developmentally regulated alternative splicing. Nat Rev Genet 12, 715–29 (2011).

3. Barbosa-Morais, N.L. et al. The evolutionary landscape of alternative splicing in vertebrate species. Science 338, 1587–93 (2012).

4. Pan, Q., Shai, O., Lee, L.J., Frey, B.J. & Blencowe, B.J. Deep surveying of alternative splicing complexity in the human transcriptome by high-throughput sequencing. Nat Genet 40, 1413–5 (2008).

5. Wang, E.T. et al. Alternative isoform regulation in human tissue transcriptomes. Nature 456, 470–6 (2008).

6. Preussner, M. et al. Body Temperature Cycles Control Rhythmic Alternative Splicing in Mammals. Mol Cell 67, 433–446 e4 (2017).

7. Fu, X.D. & Ares, M., Jr. Context-dependent control of alternative splicing by RNA-binding proteins. Nat Rev Genet 15, 689–701 (2014).

8. Martinez-Contreras, R. et al. hnRNP proteins and splicing control. Adv Exp Med Biol 623, 123–47 (2007).

9. Zhou, Z. & Fu, X.D. Regulation of splicing by SR proteins and SR protein-specific kinases. Chromosoma 122, 191–207 (2013).

10. Lee, Y. & Rio, D.C. Mechanisms and Regulation of Alternative Pre-mRNA Splicing. Annu Rev Biochem 84, 291–323 (2015).

11. Naftelberg, S., Schor, I.E., Ast, G. & Kornblihtt, A.R. Regulation of alternative splicing through coupling with transcription and chromatin structure. Annu Rev Biochem 84, 165–98 (2015).

12. Aguilera, A. & Gomez-Gonzalez, B. Genome instability: a mechanistic view of its causes and consequences. Nat Rev Genet 9, 204–17 (2008).

13. Li, X. & Manley, J.L. Cotranscriptional processes and their influence on genome stability. Genes Dev 20, 1838–47 (2006).

14. Aguilera, A. & Garcia-Muse, T. R loops: from transcription byproducts to threats to genome stability. Mol Cell 46, 115–24 (2012).

15. Aguilera, A. & Gomez-Gonzalez, B. DNA-RNA hybrids: the risks of DNA breakage during transcription. Nat Struct Mol Biol 24, 439–443 (2017).

16. Paulsen, R.D. et al. A genome-wide siRNA screen reveals diverse cellular processes and pathways that mediate genome stability. Mol Cell 35, 228–39 (2009).

17. Li, X. & Manley, J.L. Inactivation of the SR protein splicing factor ASF/SF2 results in genomic instability. Cell 122, 365–78 (2005).

18. Li, X., Niu, T. & Manley, J.L. The RNA binding protein RNPS1 alleviates ASF/SF2 depletion-induced genomic instability. RNA 13, 2108–15 (2007).

19. Xiao, R. et al. Splicing regulator SC35 is essential for genomic stability and cell proliferation during mammalian organogenesis. Mol Cell Biol 27, 5393–402 (2007).

20. Michelle, L. et al. Proteins associated with the exon junction complex also control the alternative splicing of apoptotic regulators. Mol Cell Biol 32, 954–67 (2012).

21. Yamaguchi, T. et al. p53R2-dependent pathway for DNA synthesis in a p53-regulated cell cycle checkpoint. Cancer Res 61, 8256–62 (2001).

22. Perreault, N. & Beaulieu, J.F. Use of the dissociating enzyme thermolysin to generate viable human normal intestinal epithelial cell cultures. Exp Cell Res 224, 354–64 (1996).

23. Ni, J.Z. et al. Ultraconserved elements are associated with homeostatic control of splicing regulators by alternative splicing and nonsense-mediated decay. Genes Dev 21, 708–18 (2007).

24. Irimia, M. & Roy, S.W. Origin of spliceosomal introns and alternative splicing. Cold Spring Harb Perspect Biol 6(2014).

25. Uhlen, M. et al. Proteomics. Tissue-based map of the human proteome. Science 347, 1260419 (2015).

26. O’Bryan, M.K. et al. RBM5 is a male germ cell splicing factor and is required for spermatid differentiation and male fertility. PLoS Genet 9, e1003628 (2013).

27. Sette, C., Messina, V. & Paronetto, M.P. Sam68: a new STAR in the male fertility firmament. J Androl 31, 66–74 (2010).

28. Ehrmann, I. & Elliott, D.J. Expression and functions of the star proteins Sam68 and T-STAR in mammalian spermatogenesis. Adv Exp Med Biol 693, 67–81 (2010).

29. Kress, C., Gautier-Courteille, C., Osborne, H.B., Babinet, C. & Paillard, L. Inactivation of CUG-BP1 /CELF1 causes growth, viability, and spermatogenesis defects in mice. Mol Cell Biol 27, 1146–57 (2007).

30. Hannigan, M.M., Zagore, L.L. & Licatalosi, D.D. Ptbp2 Controls an Alternative Splicing Network Required for Cell Communication during Spermatogenesis. Cell Rep 19, 2598–2612 (2017).

31. Perez, Y. et al. RSRC1 mutation affects intellect and behaviour through aberrant splicing and transcription, downregulating IGFBP3. Brain (2018).

32. Weinbauer, G., Luetjens, C., Simoni, M. & Nieschlag, E. Physiology of testicular function. in Andrology (eds. Nieschlag, E., Behre, H.M. & Nieschlag, S.) 11–59 (Springer, Berlin, Heidelberg, 2010).

33. Naro, C. et al. An Orchestrated Intron Retention Program in Meiosis Controls Timely Usage of Transcripts during Germ Cell Differentiation. Dev Cell 41, 82–93 e4 (2017).

34. Zeller, P. et al. Histone H3K9 methylation is dispensable for Caenorhabditis elegans development but suppresses RNA:DNA hybrid-associated repeat instability. Nat Genet 48, 1385–1395 (2016).

35. Zhuo, D., Madden, R., Elela, S.A. & Chabot, B. Modern origin of numerous alternatively spliced human introns from tandem arrays. Proc Natl Acad Sci U S A 104, 882–6 (2007).

